# Identification of key determinants of *Staphylococcus aureus* vaginal colonization

**DOI:** 10.1101/761841

**Authors:** Liwen Deng, Katrin Schilcher, Lindsey R. Burcham, Jakub M. Kwiecinski, Paige M. Johsnon, Steven R. Head, David E. Heinrichs, Alexander R. Horswill, Kelly S. Doran

## Abstract

*Staphylococcus aureus* is an important pathogen responsible for nosocomial and community acquired infections in humans, and methicillin-resistant *S. aureus* (MRSA) infections have continued to increase despite wide-spread preventative measures. *S. aureus* can colonize the female vaginal tract and reports have suggested an increase in MRSA infections in pregnant and postpartum women as well as outbreaks in newborn nurseries. Currently, little is known about specific factors that promote MRSA vaginal colonization and subsequent infection. To study *S. aureus* colonization of the female reproductive tract in a mammalian system, we developed a mouse model of *S. aureus* vaginal carriage and demonstrated that both hospital-associated and community-associated MRSA isolates can colonize the murine vaginal tract. Immunohistochemical analysis revealed an increase in neutrophils in the vaginal lumen during MRSA colonization. Additionally, we observed that a mutant lacking fibrinogen binding adhesins exhibited decreased persistence within the mouse vagina. To further identify novel factors that promote vaginal colonization, we performed RNA-sequencing to determine the transcriptome of MRSA growing *in vivo* during vaginal carriage at 5 hours, 1-day, and 3-days post-inoculation. Over 25% of bacterial genes were differentially regulated at all time points during colonization compared to laboratory cultures. The most highly induced genes were those involved in iron acquisition, including the Isd system and siderophore transport systems. Mutants deficient in these pathways did not persist as well during *in vivo* colonization. These results reveal that fibrinogen binding as well as the capacity to overcome host nutritional limitation are important determinants of MRSA vaginal colonization.

**IMPORTANCE:** *Staphylococcus aureus* is an opportunistic pathogen able to cause a wide variety of infections in humans. Recent reports have suggested an increasing prevalence of MRSA in pregnant and postpartum women, coinciding with the increased incidence of MRSA infections in the NICU and newborn nurseries. Vertical transmission from mothers to infants at delivery is a likely route of MRSA acquisition by the newborn, however, essentially nothing is known about host and bacterial factors that influence MRSA carriage in the vagina. Here, we established a mouse model of vaginal colonization and observed that multiple MRSA strains can persist in the vaginal tract. Additionally, we determined that MRSA interactions with fibrinogen as well as iron uptake can promote vaginal persistence. This study is the first to identify molecular mechanisms which govern vaginal colonization by MRSA, the critical initial step preceding infection and neonatal transmission.

## INTRODUCTION

*Staphylococcus aureus* is a commensal of approximately 20% of the healthy adult population (1) and also an opportunistic bacterial pathogen able to cause a wide variety of infections ranging in severity from superficial skin lesions to more serious invasive and life-threatening infections such as endocarditis and septicemia. Prevalence of *S. aureus* infections has increased due to higher rates of colonization, immunosuppressive conditions, greater use of surgical implants, and dramatic increases in antibiotic resistance (2, 3). Compared to antibiotic-susceptible strains, methicillin-resistant *S. aureus* (MRSA) infections exhibit elevated mortality rates, require longer hospital stays, and exert a higher financial burden on patients and healthcare institutions (4). Over the past 20 years, MRSA strains have expanded from healthcare settings and began infecting otherwise healthy individuals in the community (“community-associated” MRSA (CA-MRSA))(5). USA300 isolates are the most problematic lineage of CA-MRSA that have emerged and clonally expanded across the US, reaching epidemic levels in many hospital settings (6, 7).

Methicillin-susceptible *S. aureus* and MRSA possesses many virulence factors that promote bacterial persistence and invasive infections in different host sites. These virulence factors include cell wall-anchored surface proteins that facilitate *S. aureus* adherence to and invasion of host cells (8), proteases that modulate the host immune response to the bacterium (9), as well as pore-forming toxins such as α-toxin and the bicomponent leukocidins that lyse host cells (10). The expression of these various virulence determinants is dependent on factors such as growth rate, the availability of certain nutrients, host interactions, and the presence of antimicrobial compounds (8, 11-13).

Nasal carriage is known to be a risk factor for *S. aureus* infections both in the hospital and in the community with individuals often being infected with the strain that they carry (14). *S. aureus* can colonize the moist squamous epithelium in the anterior nares (15, 16), a process which depends upon specific interactions between bacterial cell adhesins and epithelial cell ligands. Two *S. aureus* surface proteins, clumping factor B (ClfB) and iron regulated surface determinant A (IsdA), have been strongly implicated in nasal colonization. Both ClfB and IsdA were shown to promote adhesion to nasal epithelium *in vitro* (17) and colonization of the nares of rodents (18, 19) and, in the case of ClfB, humans (20). ClfB is a member of a family of proteins that are structurally related to clumping factor A (ClfA), the archetypal fibrinogen (Fg) binding protein of *S. aureus*. ClfB has been shown to bind Fg, as well as cytokeratin 10, by the “dock, lock, and latch” mechanism first defined for the Fg binding proteins SdrG and ClfA (21, 22). Additional surface proteins shown to contribute to bacterial attachment to nasal epithelial cells *in vitro* include *S. aureus* surface protein G (SasG) and the serine-aspartate repeat proteins SdrC and SdrD (23).

While a ubiquitous colonizer of the skin and mucous membranes, *S. aureus*, including antibiotic sensitive and resistant strains, has also been reported to colonize the vagina in up to 22% of pregnant women (24-29). A study that examined MRSA colonization showed that out of 5,732 mothers, 3.5% were colonized by MRSA in the genital tract during pregnancy (24). Another recent study of 1834 mothers showed that 4.7% were colonized vaginally by multidrug-resistant *S. aureus* (30). Reports have suggested an increasing prevalence in the USA300 lineage of MRSA in pregnant and postpartum women, coinciding with the increased incidence in the NICU and in newborn nurseries (31-36). MRSA outbreaks in NICUs can be difficult to control and have been associated with significant morbidity and mortality (33). Vertical transmission from mothers to infants at delivery has been proposed as a possible mechanism of neonatal CA-MRSA acquisition (30, 37), and while it is clear that *S. aureus* and MRSA can colonize the vaginal tract during pregnancy, essentially nothing is known about specific bacterial factors that promote vaginal persistence.

In this study, we have adapted a murine model of vaginal colonization by Group B *Streptococcus* (GBS) (38), to investigate MRSA vaginal colonization. We determined that divergent MRSA strains, CA-MRSA USA300 and HA-MRSA252, can persist within the mouse vaginal tract and that three mouse strains, CD-1, C57BL/6, and BALB/c, can be colonized with MRSA. We detected fluorescent MRSA in the vaginal lumen as well as cervical and uterine tissues of colonized mice and immunohistochemical staining showed an increase of neutrophils in colonized mice compared to naïve mice. We found that a MRSA strain lacking fibrinogen-binding surface adhesins was attenuated in both *in vitro* and *in vivo* models of vaginal colonization. Lastly, RNA-sequencing analysis of bacteria growing *in vivo* revealed the importance of iron homeostasis in promoting MRSA persistence within the mouse vagina. Mutant USA300 strains lacking the siderophore transporter FhuCBG or the cell-surface heme receptor IsdB were significantly attenuated in their ability to colonize the vaginal tract *in vivo*.

## RESULTS

### MRSA colonization of the reproductive tract

To characterize the ability of MRSA to attach to epithelial cells of the lower female reproductive tract, we performed quantitative adherence assays with community-associated USA300 strain LAC (39) and hospital-acquired strain MRSA252 (40) as described in (41) and in the Methods. An inoculum of 10^5^ CFU/well (MOI = 1) was added to confluent monolayers of immortalized human vaginal (VK2), ectocervical (Ect1), and endocervical (End1) epithelial cells. Following a 30-minute incubation, the cells were washed to remove all nonadherent bacteria. Data are expressed as percent recovered cell-associated MRSA relative to the initial inoculum. Both strains exhibited substantial adherence to all three cell lines, ranging from 30-57% of the original inoculum (Fig. 1A and B). Next, we assessed the ability of both MRSA strains to initiate colonization of the murine vaginal tract. 8-week old female CD-1 mice were treated with 17β-estradiol 1-day before inoculation with 10^7^ CFU of either USA300 or MRSA252. The next day the vaginal lumen was swabbed and then we euthanized the animals and collected the vagina, cervix, and uterus from each mouse to quantify bacterial load. The total CFU from the swab or tissue homogenates was determined by plating on *S. aureus* CHROMagar supplemented with cefoxitin. Both strains of MRSA were recovered from the majority of mice 1-day post-inoculation in all tissues, and the CFU recovered from the swab were similar to the total CFU counts from the vaginal tissue homogenates (Fig. 1C and D). This level and range in recovered CFU is similar to what we have observed using this mouse model for GBS colonization (38). In a subsequent experiment, mice were inoculated with USA300 expressing a fluorescent DsRed protein and we harvested the female reproductive tract 1-day post-colonization for histological analysis. We made serial sections of these tissues and performed H&E staining to examine overall tissue morphology (Fig 1E, G, and I) and fluorescent microscopy to visualize USA300 (Fig 1F, H, J, K, and L). We observed numerous red fluorescent bacteria contained within the lumen of the vagina (red arrows) (Fig. 1F). We could also see MRSA in the cervical and uterine lumen, as well as within the lamina propria of those organs (green arrows) (Fig. 1H, J, K, and L)

**Figure 1.**
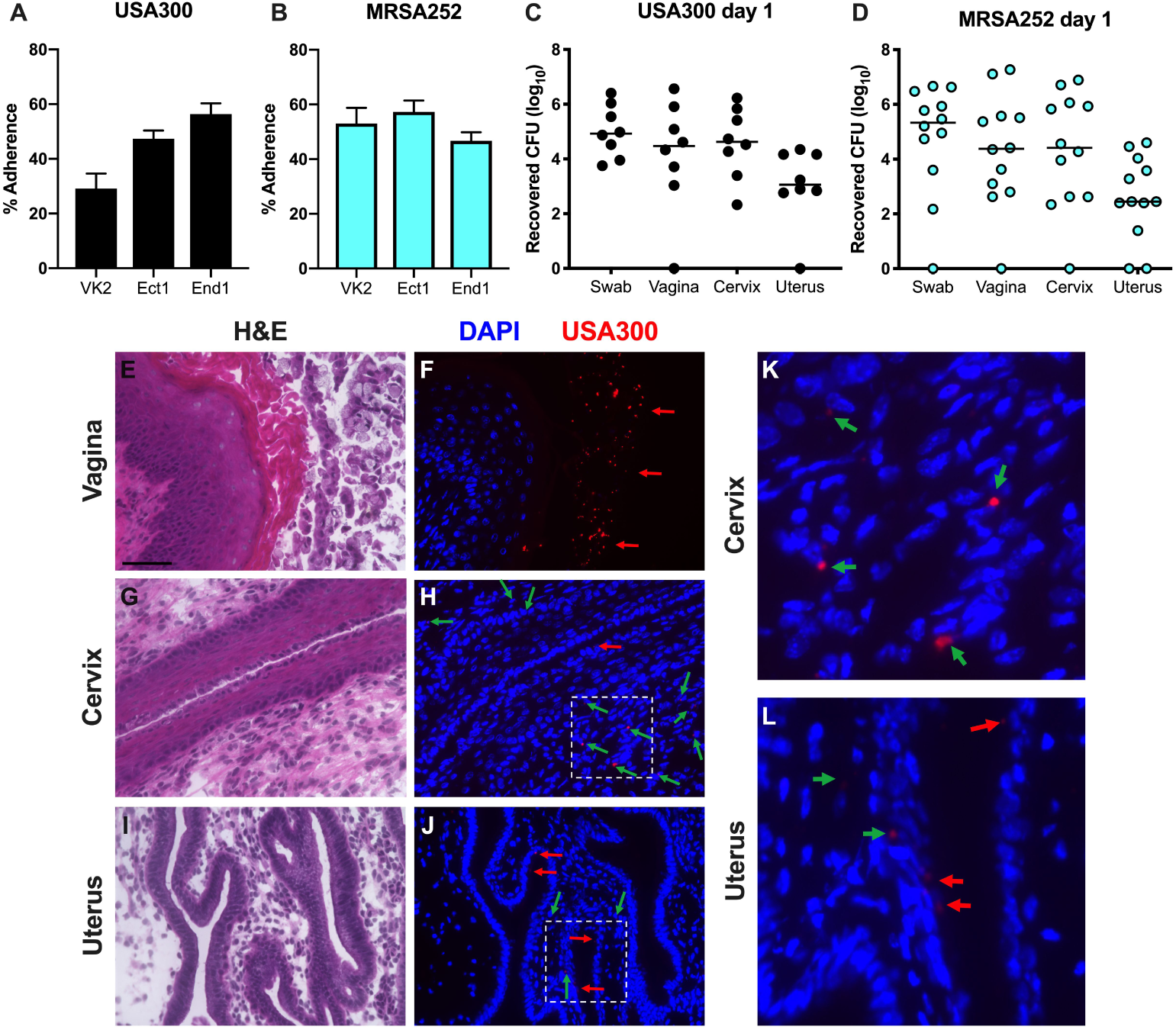
Modeling MRSA vaginal colonization. (A and B) Adherence of USA300 (A) and MRSA252 (B) to human vaginal (VK2), ectocervical (Ect1), and endocervical (End1) endothelial cells. (C and D) CFU counts from vaginal swabs, vagina, cervix, and uterus recovered 1-day post-inoculation with USA300 (C) or MRSA252 (D). Lines represent median CFU. (E-L) Mice were colonized with DsRed expressing USA300. 1-day post-inoculation, the female reproductive tract was harvested and 6μm sections of the vagina (E and F), cervix (G, H, and K), and uterus (I, J, and L) were either stained with H&E (E, G, I) or labelled with DAPI and imaged with an epifluorescent microscope to visualize nuclei and USA300 (F, H, J, K, and L). The areas highlighted in (H) and (J) are expanded in (K) and (L). USA300 in the lumen of tissues are indicated with red arrows, and USA300 within the lamina propia are indicated with green arrows. Scale bar in (E) is 100μm.

### MRSA vaginal persistence and host response

To assess vaginal persistence, mice were colonized with USA300 or MRSA252 and swabbed to determine bacterial load over time. We recovered similar CFU from mice colonized with either MRSA strain and we observed that both strains exhibited similar persistence within the mouse vagina. While all mice were initially highly colonized by both MRSA strains, some remained highly colonized while MRSA was cleared from other mice. (Fig. 2A and B). We also assessed USA300 vaginal colonization for multiple mouse strains and observed the highest mean CFUs from BALB/c mice while C57BL/6 and CD-1 mice were colonized to a lower level (Fig. S1). Furthermore, MRSA was cleared more rapidly from the vaginal tract of CD-1 mice and persisted the longest in BALB/c mice (Fig. S1).

**Figure 2.**
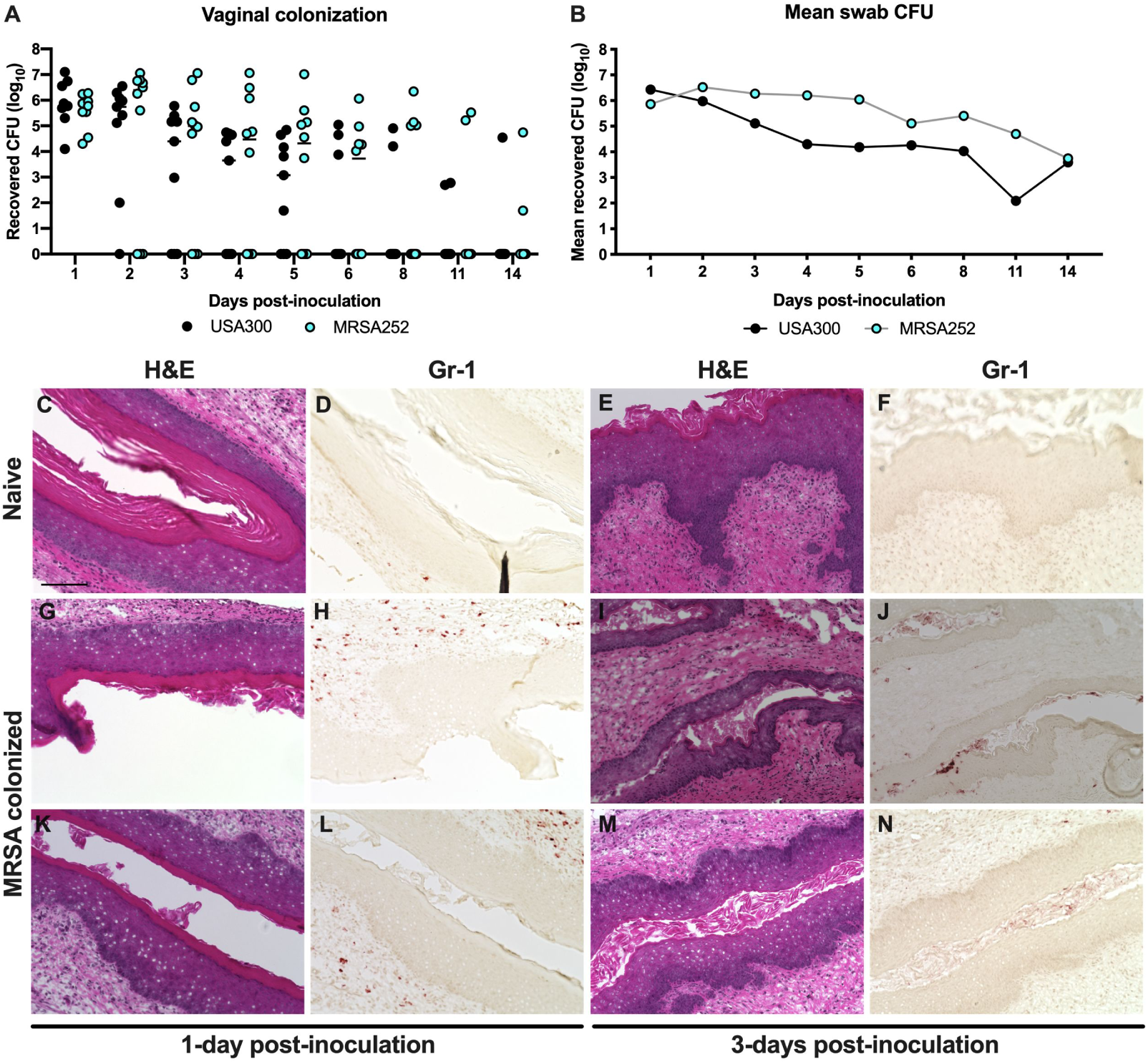
MRSA vaginal persistence and host response. (A and B) USA300 and MRSA252 persistence within the CD-1 mouse vaginal tract. CFU counts for individual mice (A) and mean recovered CFU from vaginal swabs (B) were monitored for 14 days. Lines in (A) represent median CFU. (C-N) Histology of the mouse vagina during MRSA colonization. Mice were pre-treated with 17β-estradiol and either remained naïve (C-F) or were inoculated with 10^7^ CFU of USA300 (G-N). 6μm serial sections were stained with H&E (C, E, G, I, K, M) or labelled with an antibody against Gr-1 (D, F, H, J, L, N). Scale bar in (C) is 100μm.

As we observed eventual clearance of MRSA from the vaginal tract, we examined the presence of neutrophils in vaginal tissue of mice colonized with MRSA compared to naïve mice. Previous studies have shown that neutrophils respond to vaginal colonization by pathogenic *Streptococcus* species, namely GBS and *Streptococcus pyogenes* (Group A *Streptococcus*, GAS), and that neutrophils contribute to host defense and ultimate bacterial clearance (42-44). To visualize neutrophils during colonization by MRSA, we collected vaginal tissues from mice 1-day and 3-days post-inoculation with USA300 and made serial sections for H&E staining and labelling with an antibody against the neutrophil marker Gr-1. H&E analysis showed that there were no obvious differences in morphology of the vaginal lumen between naïve and colonized mice (Fig. 2C, E, G, I, K, and M). We observed very few Gr-1 positive cells in the tissue sections form naïve mice (Fig. 2D and F). In contrast to those from naïve mice, the tissue sections from mice colonized with USA300 for 1-day contained numerous neutrophils within the vaginal lamina propria (Fig. 2H and J). At 3-days post-inoculation we detected neutrophils within the vaginal lumen (Fig. 2L and N).

### Adherence to fibrinogen impacts MRSA vaginal colonization

In a previous study, we demonstrated that GBS Fg binding contributed to vaginal persistence (45). Also, several studies have shown the importance of *S. aureus* interactions with extracellular matrix components, including Fg, in colonization and disease progression (46-49). USA300 binding to Fg is primarily mediated by the four sortase-anchored surface adhesins ClfA, ClfB, FnbA, and FnbB. (8, 46, 50). The serine-aspartate adhesins SdrC, SdrD, and SdrE are in the same protein family as ClfA/B (51) and have been reported to bind nasal epithelia (23). To eliminate these adherence functions, a USA300 strain was engineered where all of these adhesins were deleted or disrupted by incorporating four separate mutations (Δ*clfA clfB*::Tn Δ*fnbAB sdrCDE*::Tet; hereafter called “Fg adhesin mutant”). Compared to WT USA300, the Fg adhesin mutant was significantly less adherent to Fg (Fig. 3A). Quantitative adherence assays showed that the fibrinogen adhesin mutant exhibited decreased attachment to VK2 vaginal epithelial cells (Fig. 3B), and we could visualize this difference via Gram staining (Fig. 3C and D). Further, the Fg adhesin mutant was also less adherent to Ect1 and End1 cervical epithelial cells (Fig. 3E and F). To assess the impact of these important surface adhesins during *in vivo* colonization, we co-challenged mice with both WT USA300 and the Fg adhesin mutant. Initially we recovered similar CFUs of both strains from the mice. However, by 3-days post-inoculation, mice were significantly less colonized by the Fg adhesin mutant compared to WT USA300 (Fig. 3G). At 5-days post-inoculation, we could recover WT USA300 CFU from 60% of the mice while only 30% were still colonized by the Fg adhesin mutant (Fig. 3G).

**Figure 3.**
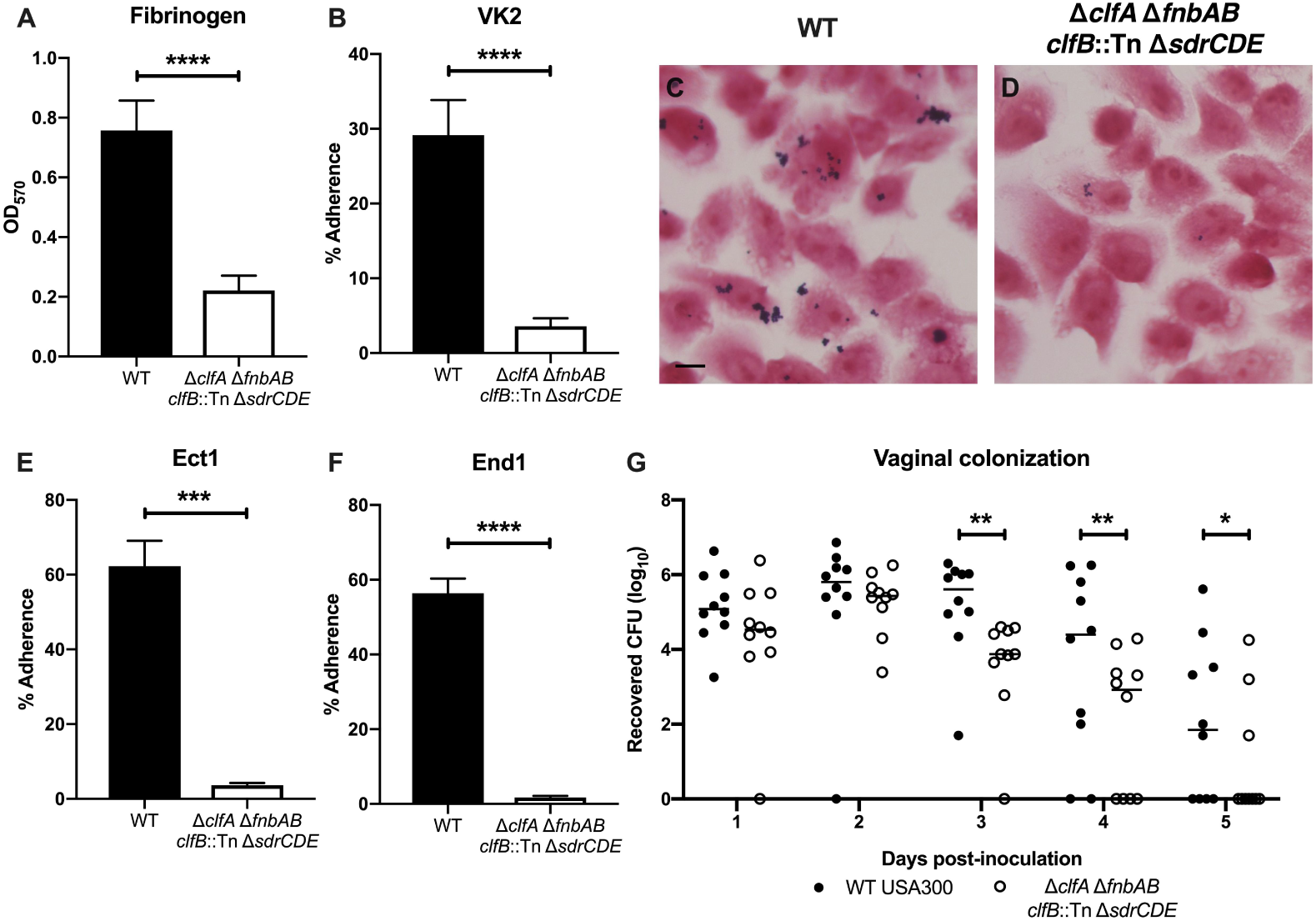
Adherence to fibrinogen impacts MRSA vaginal colonization. (A) Adherence of WT USA300 and the Fg adhesin mutant to Fg. (B-D) Adherence to VK2 cells. Monolayers of VK2 cells were inoculated with WT USA300 or the Fg adhesin mutant for a quantitative adherence assay (B) or Gram stains (C and D). Scale bar in (C) is 10 μm. (E and F) Adherence to Ect1 (E) and End1 (F) epithelial cells. (G) WT USA300 and the Fg adhesin co-colonization. Statistical analysis (A, B, E, and F) Unpaired t test. (G) Two-way ANOVA with Sidak’s multiple comparisons test. * P < 0.05; *** P < 0.0005; **** P < 0.00005.

### Transcriptome analysis during MRSA vaginal colonization

Although the Fg adhesin mutant was impaired in vaginal persistence compared to WT USA300, we did not observe a significant difference in recovered CFUs between the two strains during the first two days of colonization, and a few mice remained colonized with the Fg adhesin mutant at later time points (Fig. 1G). Thus, we hypothesized that other bacterial factors are involved in promoting MRSA vaginal carriage. To determine the impact of vaginal colonization on MRSA gene expression, we performed transcriptome analysis by RNA-sequencing of USA300 recovered from the mouse vagina compared to USA300 cultured under laboratory conditions. For these experiments, we utilized the CD-1 mouse strain as likely in this background the bacteria encounter more host pressure to maintain colonization. Mice were pre-treated with 17β-Estradiol, inoculated with 10^7^ CFU of USA300, and swabbed at 5hrs, 1-day, and 3-days post-inoculation for RNA isolation. The same mice were swabbed 2-, 4-, 6-, and 8-days post-inoculation for CFU enumeration (Fig. 4A and B). Based on swab CFU counts, we selected samples from 18 mice (purple circles) for RNA-sequencing analysis (Fig. 4B). RNA samples from 6 mouse swabs were pooled to generate 3 replicates for each time-point to compare to triplicate culture samples. Principle component analysis (PCA) for all of the samples showed that culture samples clustered separately from mouse samples (Fig. 4C). Next, we compared mouse samples from each time point to the culture samples and observed 709 genes were significantly down-regulated (Fig. 4D) and 741 genes were significantly upregulated (Fig. 4E) in the mouse (Table. S1.) Volcano plots of the log_2_(fold change) vs. –log_10_(P value) show that many of the differentially upregulated and downregulated changes were highly significant at all three time points compared to culture (Fig. 4F-H). We observed significant overlap in differentially expressed transcripts at the various time points; over half of the differentially upregulated and downregulated genes were the same at all three time points (Fig. 4E).

**Figure 4.**
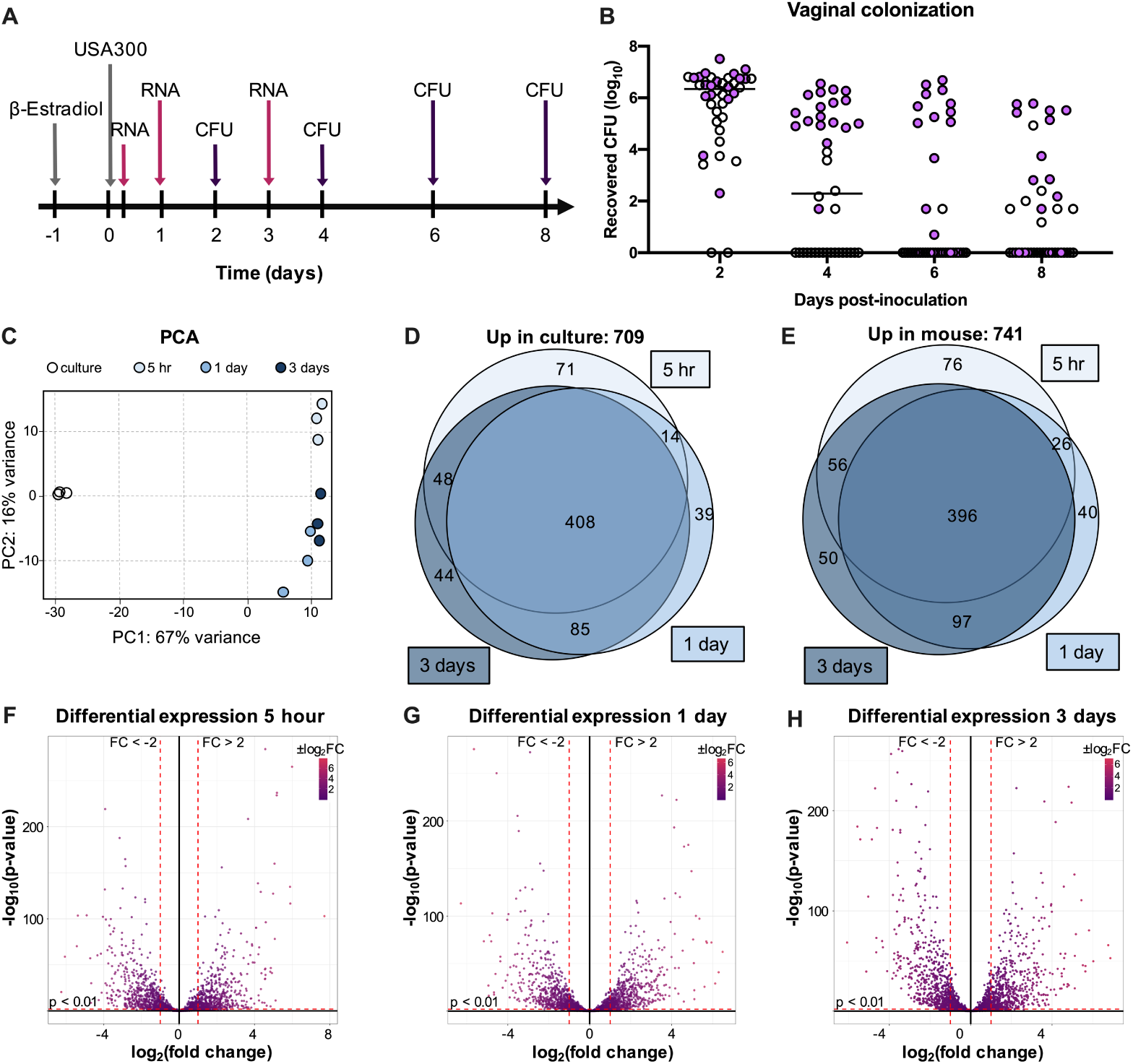
Transcriptome analysis during MRSA vaginal colonization. (A) Experimental design for RNA-sequencing analysis of mouse vaginal swabs. (B) CFU counts from mouse vaginal swabs. Samples chosen for RNA-sequencing are highlighted in purple. (C) PCA plot for triplicate samples of culture, 5hr, 1-day, and 3-days swabs. (D and E) Venn diagrams showing genes expressed at significantly higher levels (fold change > 2, P value < 0.01) in culture (D) or in mouse swab samples (E). (F-H) Volcano plots highlighting genes that are differentially expressed in swab samples from 5hrs (F), 1-day (G), and 3-days (H) post-inoculation compared to culture.

We identified genes encoding transcriptional regulators, toxins, extracellular enzymes, and extracellular matrix-binding surface proteins that were significantly upregulated at all three time points (Table 1). Interestingly, while only one immune evasion factor, chemotaxis inhibitor (*chs*), was upregulated at all three time points, additional immune evasion genes were significantly upregulated at 3-days post-inoculation. We observed a similar trend with genes encoding components of the type VII secretion system (T7SS), which has been shown to contribute to *S. aureus* virulence and competition with other microbes in polymicrobial settings (52, 53). At 5 hrs post-inoculation, only 5 T7SS genes were significantly upregulated while 14 genes were upregulated at 1-day and 3-days post-inoculation (Table 1).

**Table 1.**
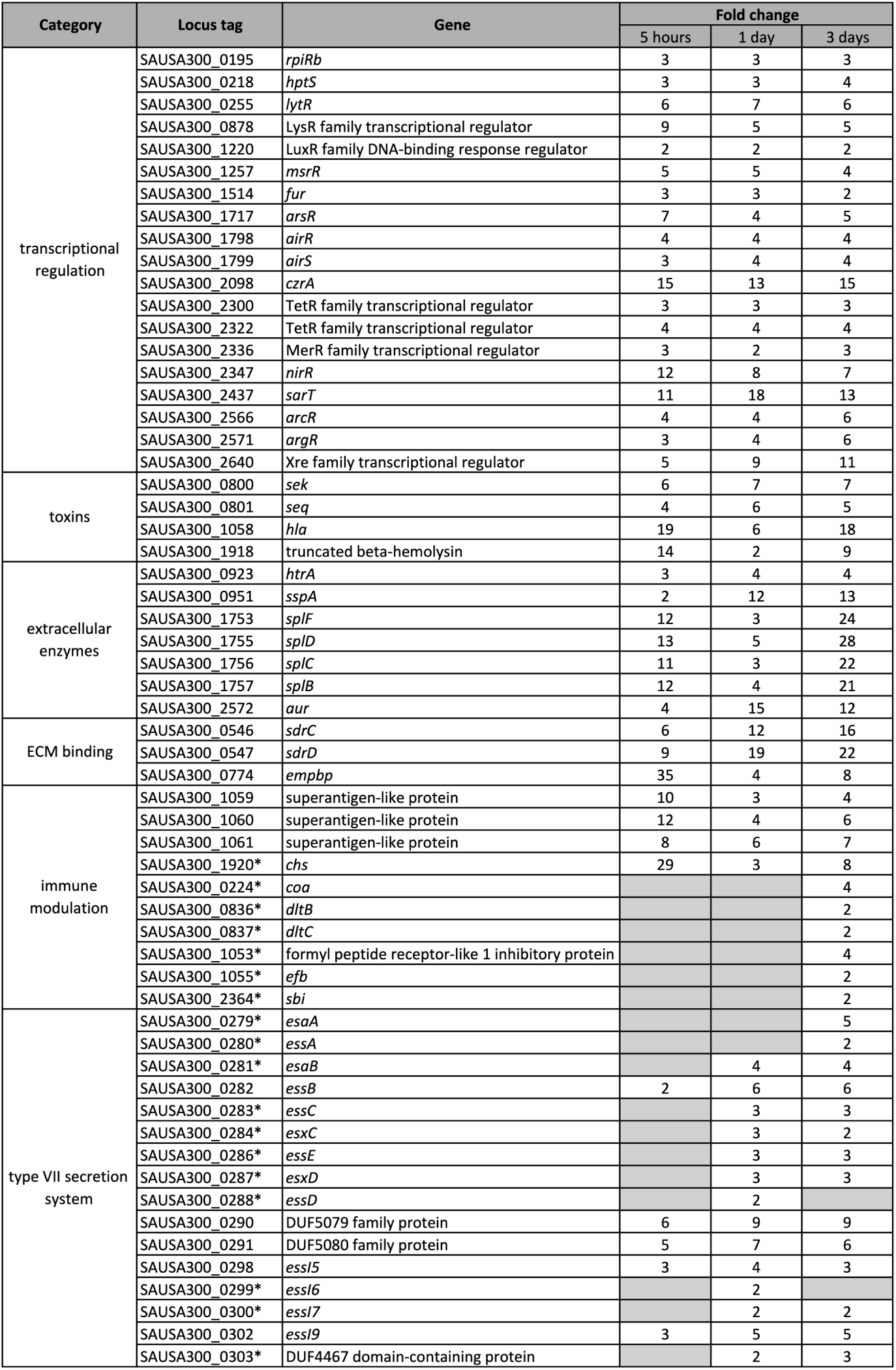
Virulence factors which were significantly upregulated during vaginal colonization. Genes which encode transcriptional regulators, toxins, secreted enzymes, and ECM binding factors and were differentially expressed in mouse swab samples at all three time points are listed with their respective fold changes relative to growth in TSB. Immune modulation and T7SS genes which were not significantly differentially expressed at all time points are indicated with asterisks.

### Iron homeostasis impacts vaginal persistence

Though there were global transcriptional changes, the most highly-significant, differentially-expressed transcripts belonged to iron uptake and export systems. The most highly induced was the iron-surface determinant *isd* heme acquisition system (*isdBACDEFG* and *srtB)*. Other genes included those involved in the production of the siderophore staphyloferrin B (SB) (*sbnABCDEFGHI*) as well as its importer (*sirAB*), the staphyloferrin A (SA) importer (*htsABC*), the xeno-siderophore transporter (*fhuCB*), as well as the catechol/catecholamine iron transporter system (*sstABCD*). Lastly, the heme-regulated export *hrt* system, was highly down-regulated during colonization (*hrtAB*). (Fig. 5A) (54-57). As these results strongly suggest that the vaginal environment is iron limited, we performed inductively coupled plasma mass spectrometry (ICP-MS) to determine the iron concentration in vaginal lavage fluid from naïve mice and mice colonized with USA300. We observed a very low concentration of iron (0.52 μM), irrespective of MRSA colonization, compared to the level present in tryptic soy broth (TSB) (10 μM) (Fig. 5B).

**Figure 5.**
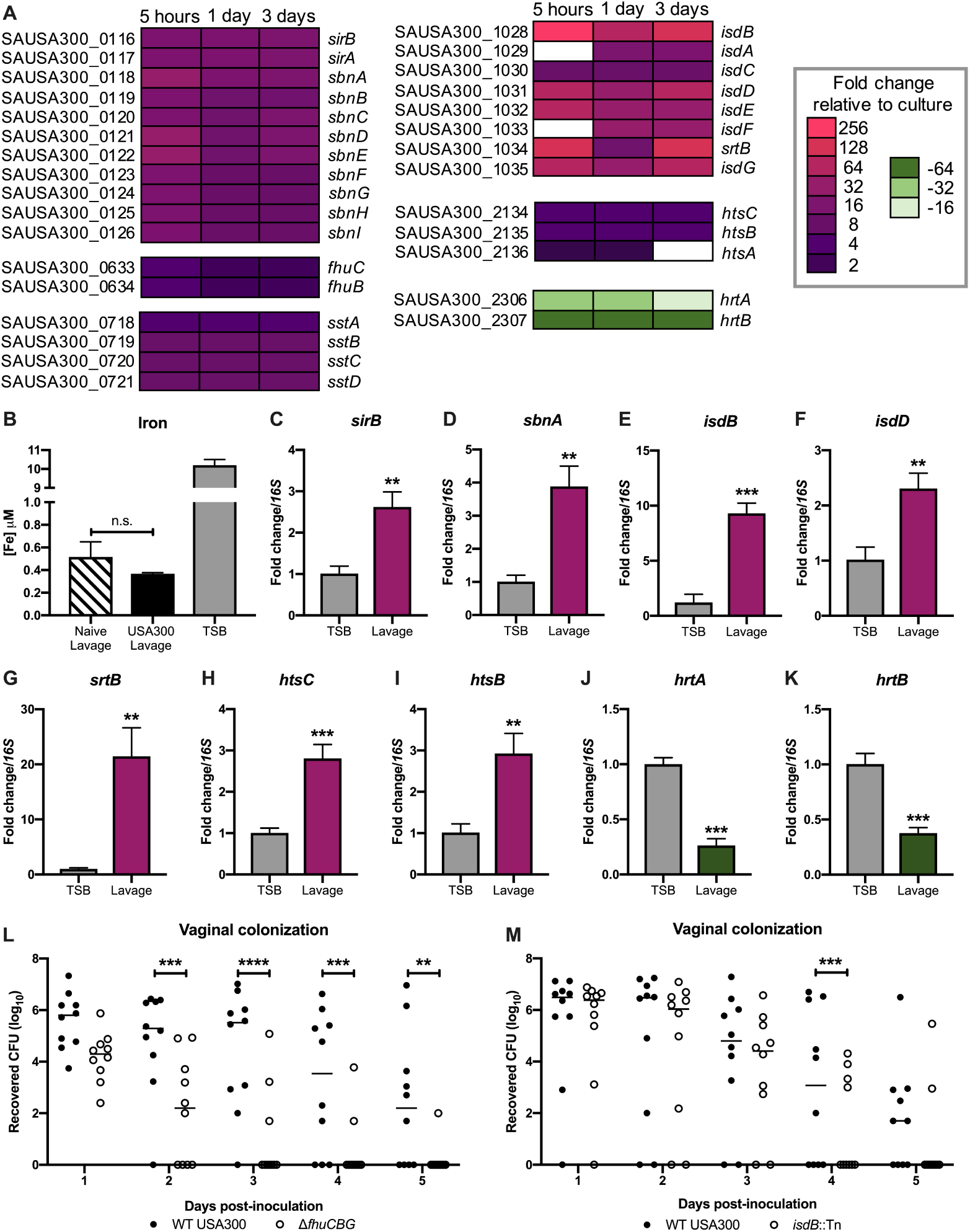
Iron homeostasis impacts vaginal persistence. (A) Differential expression of genes in iron-acquisition and iron-export pathways. (B) ICP-MS analysis of vaginal lavage from naïve and colonized mice, and TSB. (C-K) RT-qPCR confirmation of select RNA-sequencing iron-homeostasis hits. (L) Co-colonization with WT USA300 and Δ*fhuCBG* mutant. (M) Co-colonization with WT USA300 and *isdB*::Tn mutant. Statistical analysis: (B) One-way ANOVA. (C-K) Unpaired t test. (L and M) Two-way ANOVA with Sidak’s multiple comparisons test.

To confirm the differential expression of iron-uptake systems by USA300, we incubated USA300 in mouse vaginal lavage fluid and performed RT-qPCR to compare transcripts of select iron-homeostasis genes between bacteria grown in lavage fluid and bacteria cultured under laboratory conditions in TSB. Similar to our RNA-seq results, the RT-qPCR analysis revealed an increase in *sirB, sbnA, isdB, isdD, srtB, htsC*, and *htsB* transcripts in MRSA cultured in vaginal lavage fluid (Fig. 5C to 5I). Additionally, *hrtA* and *hrtB* were significantly downregulated in MRSA from vaginal lavage compared to MRSA grown in TSB (Fig. 5J and 5K).

To assess the impact of iron uptake by MRSA on vaginal persistence, we co-colonized mice with WT USA300 and either Δ*fhuCBG* or *isdB*::Tn mutants. In addition to its role in the uptake of xeno-siderophores, the FhuC ATP-ase also provides energy needed for uptake of the siderophores staphyloferrin A and staphyloferrin B. Therefore, the Δ*fhuCBG* mutant is defective in the transport of all siderophores (58). Also, our RNA-seq results show that at all three time points, the most highly upregulated gene was *isdB*, which encodes the hemoglobin-binding surface protein that transports heme to downstream components of the *isd* system (55). *isdB* transcripts from mouse samples were increased 210-fold at 5hrs, 90-fold at 1-day, and 117-fold at 3-days post-inoculation compared to culture (Table S1 and Fig. 5A). Compared to WT USA300, the Δ*fhuCBG* mutant and the *isdB*::Tn mutant were cleared significantly faster from the mouse vagina (Fig. 5L and M). Because a previous study reported that IsdB may impact bacterial attachment to host cells (59), we quantified adherence of the *isdB*::Tn mutant to VK2, Ect1, and End1 cells *in vitro* and observed no defect compared to WT USA300 (Fig. S2).

## DISCUSSION

*S. aureus* is capable of causing disease at nearly every site of the body (60), and MRSA colonization of the skin as well as mucosal sites, such as the nares and the vaginal tract, is a necessary initial step preceding the development of invasive disease (27, 37, 61-63). While many studies have investigated host and bacterial determinants of *S. aureus* colonization of the skin and nares as well as subsequent infection, little is known about factors which influence vaginal niche establishment and persistence. Because vaginal carriage during pregnancy represents a major risk factor for transmission of this pathogen to the newborn (24, 25, 64, 65), we utilized *in vitro* and *in vivo* models of MRSA vaginal colonization to identify determinants of persistence within the female reproductive tract. The results of our study reveal that MRSA can interact directly with the female reproductive tract epithelium *in vitro* and *in vivo*, and that the expression of cell-wall anchored Fg binding adhesins as well as iron-acquisition systems promote MRSA vaginal colonization.

The effect of *S. aureus* colonization on the host immune response has been well-characterized at many epithelial sites. *S. aureus* on the skin promotes a robust inflammatory response involving both the innate and adaptive immune system (66, 67). Neutrophils in particular are rapidly and highly recruited to the site of *S. aureus* skin infection and are key mediators of clearance of the pathogen (68-75). Our studies on GBS vaginal carriage have shown a clear role for neutrophils in combatting GBS colonization of this host site (43, 76). Additionally, neutrophils have been shown to respond to other common pathogens of the vaginal tract such as the fungus *Candida albicans* (77, 78) and the Gram-negative bacterium *Neisseria gonorrhoeae* (79). In this study, we observed an increased neutrophil presence in the vaginal tissues from mice colonized by MRSA compared to naïve controls. Interestingly, while there is obvious neutrophil infiltration of the lamina propria of the vagina 1-day post-colonization with MRSA, we did not detect neutrophils in the vaginal lumen at this early time point. In contrast, at 3-days post-inoculation, we could visualize many neutrophils within the vaginal lumen. The timing of the infiltration of neutrophils into the vaginal lumen coincides with the increased expression of immune evasion factors by MRSA; in our RNA-sequencing analysis we observed significant upregulation of these factors at 3-days post-inoculation and not at earlier time points. Future studies aimed at further characterizing the dynamics of neutrophil response to MRSA in the female reproductive tract and their extravasation into the vaginal lumen may reveal new insights into host immune responses common to all vaginal pathogens as well as those specific to MRSA.

The impact of *S. aureus* interactions with Fg on colonization and disease at various tissue sites has been well-characterized. In the context of invasive infections, Fg and fibrin can promote clearance of *S. aureus* by containing the bacteria within aggregates (80, 81). Additionally, Fg can stimulate the production of inflammatory cytokines and activate neutrophils (82-84). However, *S. aureus* has also been shown to target Fg to promote persistence and disease in the host. The bacterium can interact with Fg in order to coagulate or to form clumps which help it evade immune detection, and this clumping is mediated by surface Fg binding adhesins including ClfA, ClfB (46, 85-88). There is also evidence that *S. aureus* can alter gene regulation in the presence of fibrinogen-containing clumps to enhance expression of virulence determinants (89). Moreover, *S. aureus* can use Fg as part of its biofilm structure to promote persistence within the host (90). Our data suggest that, in the context of vaginal colonization, MRSA interactions with Fg are necessary for persistence within the host. A mutant deficient in Fg binding was significantly impaired in its ability to adhere to human female reproductive tract cells *in vitro* and was also rapidly cleared from the vaginal tract *in vivo* compared to the WT. These results hint that the benefits of MRSA binding to Fg outweigh the potential detriments for the pathogen during vaginal colonization.

While a majority of mice rapidly clear the Fg adhesin mutant during vaginal colonization, it is able to persist in some of the animals (Fig. 3G). This result suggests that there are likely other factors that contribute to *in vivo* vaginal colonization. To identify additional determinants of vaginal persistence, we performed RNA-sequencing to profile the transcriptome of MRSA during vaginal colonization. We observed that over one-quarter of the genes of USA300 were differentially expressed during *in vivo* colonization, and over half of those genes were differentially expressed at all three *in vivo* time points that were analyzed. Of note, many of the most highly and significantly differentially expressed genes belonged to iron-acquisition or iron-homeostasis pathways. Our observation that genes involved in iron uptake were upregulated was not surprising since their expression is controlled by iron levels and our ICP-MS data revealed the vaginal environment to be limited in iron (Fig. 5B). Using our *in vivo* murine vaginal colonization model, we confirmed that mutants in *fhuCBG* and *isdB* exhibited decreased persistence compared to the isogenic WT MRSA strain. Numerous reports have demonstrated the importance of nutrient iron for *S. aureus* growth and pathogenicity (55, 91, 92), and the results of our study highlight the necessity of this metal for MRSA persistence within the vaginal environment. That the Δ*fhuCBG* mutant was attenuated in this model was interesting because, while FhuCBG is known to transport hydroxamate-type siderophores which *S. aureus* does not synthesize (57, 93), FhuC is also the ATP-ase which provides energy for uptake of both SA and SB siderophores (55, 58). Both WT and the Δ*fhuCBG* mutant should, under the iron-restricted conditions during vaginal colonization, express SA and SB. Given that the Δ*fhuCBG* mutant cannot transport these siderophores, the extracellular environment becomes more iron restricted to the mutant as it cannot access SA-Fe and SB-Fe chelates.

The limitation of iron is a major host mechanism for defending against pathogens because this metal is vital for bacterial growth and metabolic processes (55, 94, 95). Other transcriptomic studies examining *S. aureus* growing *in vivo* during invasive infections have shown that the bacteria respond to nutrient limitation within the host. One study which compared the transcriptomes of *S. aureus* in a murine osteomyelitis model to bacteria grown under laboratory conditions revealed the importance of iron homeostasis mechanisms, especially the Isd pathway, during chronic infection (96). Another analysis of USA300 gene expression during human and mouse infections also showed upregulation of iron transporters *in vivo* (97). Interestingly, many reports have shown that neutrophils can play an active role in limiting iron in numerous host sites, including the vagina, during exposure to a bacterial pathogen (98-101). The precise mechanisms by which the host restricts iron availability during colonization warrants further research as this would provide insight into the exact function of neutrophils in controlling MRSA vaginal persistence.

We have developed a murine model of *S. aureus* vaginal colonization and this study is the first to investigate the molecular mechanisms that promote vaginal carriage and persistence by MRSA. This mouse model will be useful for continued studies on MRSA-host interactions within a mucosal environment. Here we demonstrate the importance of Fg binding as well as iron-acquisition in promoting long-term colonization. Additionally, we observed that neutrophils respond to MRSA presence in the vagina and that the bacteria upregulate the expression of immune-modulating genes during the course of colonization. Further investigation into these specific colonization determinants could yield therapeutic interventions to treat MRSA persistence within this host niche.

## MATERIALS AND METHODS

### Bacterial strains and culture conditions

*S. aureus* strains USA300 (39) and MRSA252 (40) were used for the experiments. *S. aureus* was grown in tryptic soy broth (TSB) at 37 °C and growth was monitored by measuring the optical density at 600nm (OD_600_). For selection of *S. aureus* mutants, TSA (tryptic soy agar) was supplemented with chloramphenicol (Cm) (10 μg/mL), erythromycin (Erm) (3 μg/mL), or tetracycline (Tet) (1 μg/mL).

To generate the Fg adhesin mutant, first the *fnbAB* operon was deleted using allelic replacement. Phages 80α or 11 were used for transduction between *S. aureus* strains (102). The *fnbAB* markerless deletion plasmid pHC94 was constructed using Gibson assembly with the plasmid backbone coming from amplification of pJB38 (103) using primers pJB R2 and pJB38 F2. The region upstream of *fnbA* was amplified with primers fnbAB delA and fnbAB delB, and the region downstream of *fnbB* was amplified using fnbAB delC and fnb delD (Table S2). The resulting plasmid was electroporated in *S. aureus* RN4220 (104), selecting on TSA Cm plates at 30°C. The plasmid was then transduced into *S. aureus* strain LAC Δ*clfA* (85). Individual colonies were streaked on TSA Cm plates incubated at 42°C to select for integration of the plasmid into the chromosome. Single colonies were grown in TSB at 30°C and re-inoculated into fresh media for several days before plating on TSA containing anhydrotetracycline (0.3 μg/mL) to select for loss of the plasmid, creating the LAC Δ*clfA* Δ*fnbAB* mutant. The *clfB*::Tn mutation was than transduced into this background from the Nebraska Transposon Mutant Library (105) and selected on TSA Erm plates.

The *isdB mariner*-based transposon *bursa aurealis* mutation (JE2 *isdB*::ΦNΣ, NE1102) from the Nebraska Transposon Library (105) into USA300 LAC with phage 11 as described previously (106). *S. aureus* genomic DNA of LAC* *isdB*::ΦNΣ was isolated using Puregene DNA purification kit (Qiagen) and the transposon insertion was verified by PCR with primers KAS249 and KAS250 (Table S2).

The *fhuCBG* mutant (58) and the DsRed expressing USA300 strain were generated previously (107).

### *In vitro* MRSA adherence assays

Immortalized VK2 human vaginal epithelial cells, Ect1 human ectocevical endothelial cells, and End1 human endocervical epithelial cells were obtained from the American Type Culture Collection (VK2.E6E7, ATCC CRL-2616; Ect1/E6E7, ATCC CRL-2614; End1/E6E7, ATCC CRL-2615) and were maintained in keratinocyte serum-free medium (KSFM; Gibco) with 0.1 ng/mL human recombinant epidermal growth factor (EGF; Gibco) and 0.05 mg/ml bovine pituitary extract (Gibco) at 37°C with 5% CO_2_.

Assays to determine cell surface-adherent MRSA were performed as described previously (41). Briefly, bacteria were grown to mid-log phase to infect cell monolayers (multiplicity of infection [MOI] = 1). After a 30-min. incubation, cells were detached with 0.1 mL of 0.25% trypsin-EDTA solution and lysed with addition of 0.4 mL of 0.025% TritonX-100 by vigorous pipetting. The lysates were then serially diluted and plated on TSA to enumerate the bacterial CFU. Experiments were performed at least three times with each condition in triplicate, and results from a representative experiment are shown.

Crystal violet fibrinogen adhesion assays were performed as described in (85). Briefly, 96-well plates (Corning) were coated with 20 μg/mL of human fibrinogen and incubated with 100 μL of bacterial suspensions in PBS at OD_600_ = 1.0 for 1h at 37°C. Wells were then washed and dried, and the adherent bacteria were stained with 0.1% crystal violet. The bound crystal violet stain was solubilized with 33% acetic acid and measured at OD_570_.

For Gram staining analysis, VK2 monolayers were grown in tissue culture treated chamber slides (ThermoFisher) and infected with either WT USA300 or the fibrinogen adhesin mutant at an MOI of 20. Following a 30 min incubation, the cell monolayers were washed to remove any non-adherent bacteria then fixed with 10% formalin (Fisher) and Gram stained (Sigma).

### Murine vaginal colonization model

Animal experiments were approved by the Institutional Animal Care and Use Committee at the University of Colorado-Anschutz Medical Campus protocol #00316 and performed using accepted veterinary standards. A mouse vaginal colonization model for GBS was adapted for our studies (38). 8-week old female CD-1 (Charles River), C57BL/6 (Jackson), and BALB/c (Jackson) mice were injected intraperitoneally with 0.5 mg 17β-estradiol (Sigma) 1 day prior to colonization with MRSA. Mice were vaginally inoculated with 10^7^ CFU of MRSA in 10μL PBS and on subsequent days the vaginal lumen was swabbed with a sterile ultrafine swab (Puritan). To assess tissue CFU, mice were euthanized according to approved veterinary protocols and the female reproductive tract tissues were placed into 500μL PBS and bead beat for 2 min to homogenize the tissues. The recovered MRSA was serially diluted and enumerated on CHROMagar (Hardy Diagnostics) supplemented with 5.2 μg/mL of cefoxitin.

### Histology

Mouse female reproductive tract was harvested and embedded into OCT compound (Sakura) and sectioned with a CM1950 freezing cryostat (Leica). For fluorescence microscopy, coverslips were mounted with VECTASHIELD mounting medium with DAPI (Vector Labs). H&E staining was performed using reagents from Sigma. Immunohistochemical analysis was performed using a biotinylated primary antibody against Gr-1 (Biolegend), Streptavidin conjugated to horse radish peroxidase (Jackson Immunoresearch), and AEC peroxidase substrate kit (Vector Labs). Images were taken with a BZ-X710 microscope (Keyence).

### Generation of RNA-sequencing data

10^7^ CFU of USA300 were inoculated into the mouse vagina and mice were swabbed vaginally 5 hrs, 1-day, and 3-days post-inoculation for RNA recovery. Vaginal swabs were placed into TRIzol reagent (Thermo Fisher), vortexed to dissociated bacteria from swabs, and stored at −80°C. Swabs samples from 6 mice were pooled and bacteria were lysed by beating for 2 min at maximum speed on a bead beater (BioSpec Products). RNA was isolated by following the manufacturer’s protocol using a Direct-Zol RNA MiniPrep Plus kit (Zymo Research). For each sample, 120 ng total RNA was ribodepleted using the Ribo-Zero Magenetic Gold Kit (Epidemiology) from Epicentre (Illumina) following the manufacurer’s protocol. Ribodeplete RNA was then prepared into sequence libraries using the RNA Ultra II kit (New Enlgand Biolabs) following the manufacturer’s protocol without fragmentation. Libraries underwent 9 cycles of PCR efore 1X Ampure Bead purification (Beckman Coulter). Libraries were quantified, pooled, and sequenced on an Illumina NextSeq500 with 75-base single reads targeting 20M reads per samples.

### Analysis of RNA-sequencing data

Sequencing reads were aligned to the NCBI reference sequence with GenBank accession number NC_007793.1 and expression levels were calculated using Geneious 11.1.5. Transcripts with an adjusted P value < 0.05 and log_2_fold change ±1 were considered significantly differentially expressed. PCA and volcano plots were generated using the ggplot2 package in R. Venn diagrams were generated using the area-proportional Venn diagram tool (BioInfoRx).

### ICP-MS analysis

Triplicate samples of mouse vaginal lavage and TSB samples were sent to the University of Nebraska Spectroscopy and Biophysics Core. Fe56 and Fe57 isotope measurements were combined to show total iron levels.

### RT-qPCR confirmation of RNA-sequencing

Vaginal lavage fluid was collected as described in (38) and filtered through 0.22μm Spin-X centrifuge tube filters (Costar) to remove contaminants. Triplicated log phase cultures of USA300 were pelleted and resuspended in filtered lavage fluid. Following a 2-hour incubation at 37°C, bacteria were collected by centrifugation and resuspended in Trizol, lysed by bead beating, and RNA was isolated using the Direct-Zol RNA MiniPrep Plus kit as described above. RNA was treated with Turbo DNase (Invitrogen) to remove contaminating DNA. cDNA was generated using the Quanta cDNA synthesis kit (Quanta Biosciences) and qPCR was performed using PerfeCTa SYBR Green reagent (Quanta) and a CFX96 Real-Time PCR thermal cycler (Bio-rad). Fold changes were calculated using the Livak method (108).

### Data analysis

GraphPad Prism version 7.0 was used for statistical analysis and statistical significance was accepted at P values of < 0.05 (* P < 0.05; ** P < 0.00005; *** P < 0.0005; **** P < 0.00005). Specific tests are indicated in figure legends.

## ACKNOWLEGMENTS

We thank Dr. Heidi A. Crosby for help in strain construction and the Array and NGS Core Facility at The Scripps Research Institute, Director Steven Head, for performing RNAseq analysis. This study was supported by the Rees Stealy Research Foundation/SDSU Heart Institute and San Diego Chapter ARCS Scholarships to L.D., an American Heart Association postdoctoral fellowship 17POST33670580 to J.M.K., the Canadian Institutes of Health Research to D.E.H., and the NIH/NIAID R21 AI130857 to A.R.H. and K.S.D.

---

**Supplemental Figure 1. USA300 vaginal colonization in different mouse strains.**

CD-1, C57BL/6, and BALB/c mice were inoculated with 10^7^ CFU of USA300 and the percent of mice colonized (A) as well as mean vaginal swab CFU (B) were monitored for 9 days.

**Supplemental Figure 2. WT USA300 and *isdB*::Tn adherence to (A) VK2, (B) End1, and (C) Ect1 cells.**

**Supplemental table 1. Differentially expressed genes in mouse swabs.** All genes that were significantly differentially expressed in mouse samples are listed along with their adjusted P values, fold changes relative to culture in TSB, and absolute confidence.

**Supplemental table 2.** List of primers used in this study.

